# Handcuffing intrinsically disordered regions in Mlh1-Pms1 disrupts mismatch repair

**DOI:** 10.1101/2021.03.02.433678

**Authors:** Christopher M. Furman, Ting-Yi Wang, Qiuye Zhao, Kumar Yugandhar, Haiyuan Yu, Eric Alani

## Abstract

The DNA mismatch repair (MMR) factor Mlh1-Pms1 contains long intrinsically disordered regions (IDRs). While essential for MMR, their exact functions remain elusive. We performed cross-linking mass spectrometry to identify the major interactions within the Mlh1-Pms1 heterodimer and used this information to insert FRB and FKBP dimerization domains into the IDRs of Mlh1 and Pms1. Yeast bearing these constructs were grown with rapamycin to induce dimerization. Strains containing FRB and FKBP domains in the Mlh1 IDR displayed complete MMR defects when grown with rapamycin, but removing rapamycin restored MMR functions. Furthermore, linking the Mlh1 and Pms1 IDRs through FRB-FKBP dimerization disrupted Mlh1-Pms1 binding to DNA, inappropriately activated Mlh1-Pms1, and caused MMR defects *in vivo*. We conclude that dynamic and coordinated rearrangements of the MLH IDRs regulate how the complex clamps DNA to catalyze MMR. The application of the FRB-FKBP dimerization system to interrogate *in vivo* functions of a critical repair complex will be useful for probing IDRs in diverse enzymes and to probe transient loss of MMR on demand.

## Introduction

Intrinsically disordered regions (IDRs), characterized as being conformationally flexible, are present in roughly 30% of eukaryotic proteins. While many IDRs have unknown roles, some facilitate interactions with other proteins or between regions of the same protein. IDRs are often critical for overall protein function, but the mechanisms of action are lacking because IDRs do not have a clear structure and amino acid substitutions in IDRs often have no effect (Oldfield & Dunker, 2014; Van Der Lee *et al*, 2014; Uversky, 2019). Here, we investigate the roles that the IDRs of the highly conserved MutL family proteins play during DNA mismatch repair (MMR).

MMR acts to correct DNA misincorporation errors that arise during replication (Kunkel & Erie, 2015). In baker’s yeast, DNA mismatches are recognized by the MutS homolog (MSH) proteins Msh2-Msh6 and Msh2-Msh3. These proteins act as ATP-modulated sliding clamps to recruit MLH proteins, principally Mlh1-Pms1, which nick the newly replicated strand of DNA in the vicinity of the mismatch through steps that require interactions with the DNA polymerase processivity clamp PCNA (Fig 1A; Gradia *et al*, 1999; Gorman *et al*, 2012; Pluciennik *et al*, 2010; Kawasoe *et al*, 2016). ATP binding by Mlh1-Pms1 stimulates its endonuclease activity on double-stranded DNA, and this stimulation is significantly greater on DNA substrates loaded with PCNA (Kim *et al*, 2019; Kadyrov *et al*, 2006; Tran & Liskay, 2000; Räschle *et al*, 2002; Genschel *et al*, 2017). Nicks located 5’ to the mismatch act as entry sites for Exo1, a 5’ to 3’ exonuclease, to digest the DNA strand containing the misincorporation error. The single stranded binding protein RPA then coats the ssDNA gap, after which a DNA polymerase (δ or ε) resynthesizes the gapped DNA. A partially redundant Exo1-independent mechanism requires Mlh1-Pms1 to make multiple nicks on the newly replicated strand in the vicinity of the mismatch, enabling Polymerases δ and/or ε to remove the mismatch via DNA synthesis and strand displacement. The nicks generated by both pathways are sealed by DNA ligase (Kadyrov *et al*, 2009; Goellner *et al*, 2014; Goellner *et al*, 2015; Hermans *et al*, 2016; Kim *et al*, 2019).

**Figure 1.**
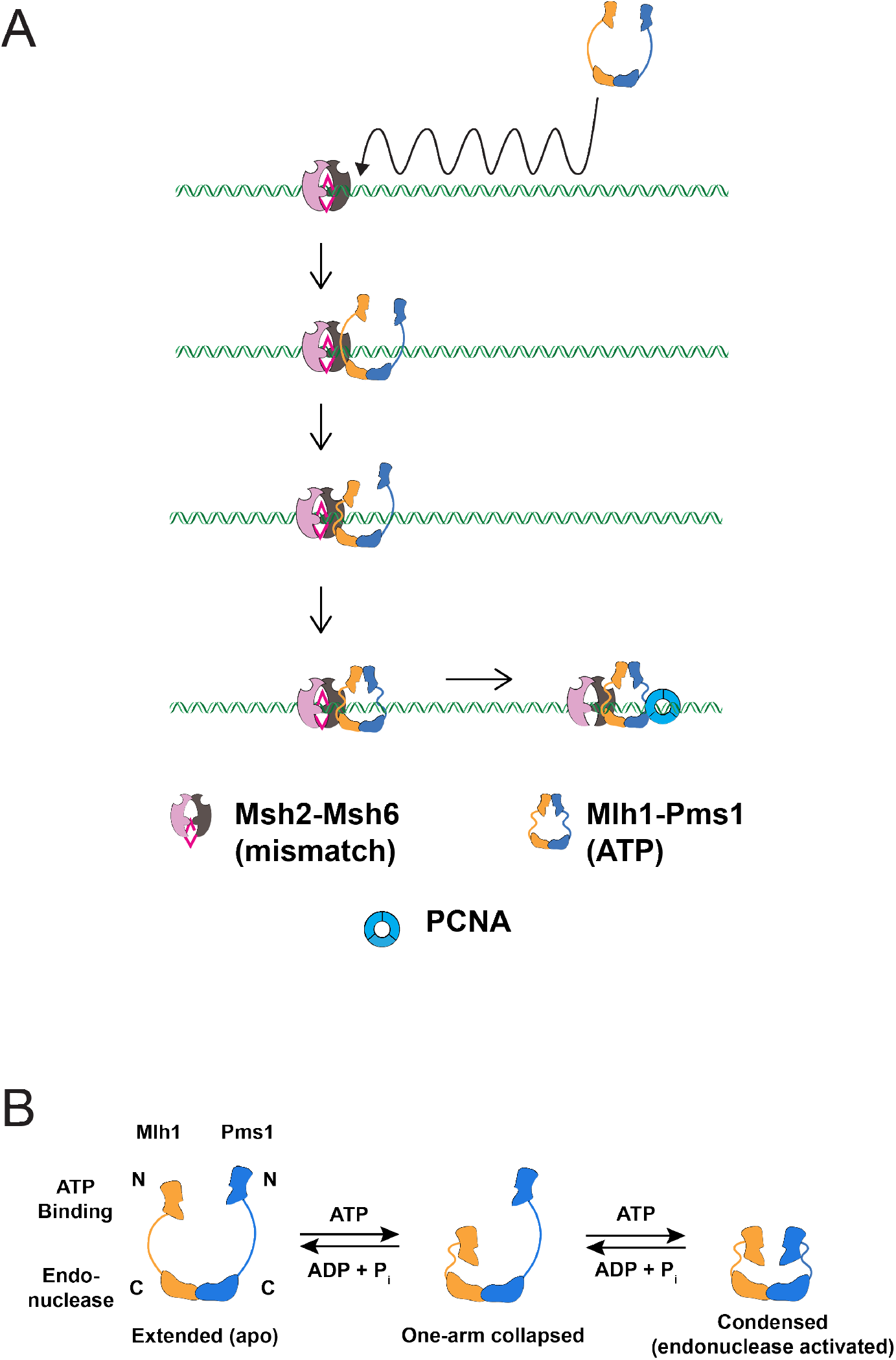
ATP-driven conformational changes in Mlh1-Pms1 during MMR. (A) A model for Mlh1-Pms1 interactions with MSH proteins during MMR. See Introduction for details. (B) Sacho *et al* (2008) proposed that ATP binding to one subunit of the MLH heterodimer promotes the formation of a one-armed collapsed state. Subsequent binding to the second subunit condenses its linker arm to yield a condensed complex that is thought to represent the activated endonuclease state.

Single molecule studies have provided a mechanistic view of how Mlh1-Pms1 interacts with MSH proteins and mismatched DNA (Fig 1A). Studies by Gorman *et al* (2010, 2012) showed that Mlh1-Pms1 is targeted to lesion-bound Msh2-Msh6 by one-dimensional hopping and three-dimensional diffusion mechanisms, and the addition of ATP provoked the release of Msh2-Msh6/Mlh1-Pms1 from mismatched DNA. How do the MLH proteins accomplish these diffusion steps? Initial hints came from atomic force microscopy and ATP hydrolysis analyses which showed that ATP binding by Mlh1-Pms1 is accompanied by conformational rearrangements involving IDRs in both subunits (Fig 1B; Hall *et al*, 2003; Sacho *et al*, 2008; Hall *et al*, 2001). Mlh1 (164 amino acids in length) and Pms1 (295 amino acids in length) each contain IDRs separated by globular N- and C-terminal domains (Appendix Fig S1A; Plys *et al*, 2012; Hall *et al*, 2003; Prolla *et al*, 1994; Argueso *et al*, 2003; Guarné *et al*, 2004). Kim *et al* (2019) showed that an MMR defective Mlh1-Pms1 complex containing shorter IDRs retained wild-type DNA binding affinity but showed diffusion defects on both naked and nucleosome-coated DNA. They also showed that the IDRs regulated the ATP hydrolysis and nuclease activities encoded by the N- and C-terminal domains of the complex, respectively. These studies suggest multi-functional roles for IDRs in regulating Mlh1-Pms1 diffusion on DNA and nucleolytic processing. Liu *et al* (2016) showed that *E. coli* MutS sliding clamps recruited MutL on mismatched DNA to form a MutS-MutL search complex. In their model they hypothesized that “an open conformation of EcMutL is required to interact with EcMutS sliding clamps, which then binds ATP to form a second, exceedingly stable, ring-like clamp” that acts as a search complex (Liu *et al*, 2016; Groothuizen *et al*, 2015).

The above models argue that the MLH IDRs play key roles in facilitating these steps, but it remains unclear how they occur (Kim *et al*, 2019; Sacho *et al*, 2008, Mardenborough *et al*, 2019). Sacho *et al* (2008) and Kunkel and Erie (2015) proposed that ATP binding to the Mlh1 subunit of the Mlh1-Pms1 heterodimer promotes the formation of a one-armed collapsed complex in which the N-terminal ATP binding domain of Mlh1 and its IDR fold into or near the C-terminal Mlh1-Pms1 domain (Fig 1B). ATP binding to Pms1 then condenses its linker arm to yield a condensed complex that positions the N-terminus of Mlh1 and Pms1 in proximity to the C-terminal domains. This conformational change, modulated through the IDR domains of both proteins, is hypothesized to change Mlh1-Pms1 affinity for DNA and activate its endonuclease activity located at the C-terminus of Pms1 (Tran & Liskay, 2000; Sacho *et al*, 2008; Ban *et al*, 1999; Pillon *et al*, 2010; Gueneau *et al*, 2013).

We performed a direct test of the above models by inserting FRB and FKBP dimerization domains into the Mlh1 and Pms1 IDRs. The FRB and FKBP domains form tight dimers upon the addition of the small molecule rapamycin (Spencer *et al*, 1993). Introducing both FRB and FKBP domains into the Mlh1 IDR conferred a null MMR phenotype in the presence of rapamycin, showing that changes in conformation in the Mlh1 IDR are critical for MMR. We then inserted FRB and FKBP domains into each IDR and showed that strains containing these constructs displayed strong MMR defects when grown in the presence of rapamycin. *In vitro* studies of one such complex showed that MMR defects were accompanied by the inappropriate activation of Mlh1-Pms1 and a defect in Mlh1-Pms1 binding to DNA. Our observations indicate that the MLH IDRs mediate dynamic conformational steps required for Mlh1-Pms1 to form a clamp on DNA and for licensing subsequent repair steps. Importantly, we provide a general strategy to reversibly lock conformational changes in IDR domains that can be used to study a diverse number of cellular processes.

## Results

### Mlh1-Pms1 inter-subunit interactions involve the N-terminal, IDR and C-terminal domains of both proteins

We used cleavable cross-linking coupled with mass spectrometry (XL-MS) as a tool to map intra- and inter-subunit interactions in Mlh1-Pms1. In this approach, disuccinimidyl sulfoxide (DSSO; Materials and Methods) was used to cross-link exposed lysine residues in close proximity (10.1 angstrom spacer length; Yu & Huang, 2018; Yugandhar *et al*, 2020). While this method cannot yield detailed structural information, it can identify regions of proteins that potentially interact and thus give insights into the movements of Mlh1-Pms1 during MMR. We were unable to perform XL-MS in the presence of ATP or the non-hydrolysable ATP analog AMP-PMP because the primary amines present in these molecules interfered with our detection of DSSO-induced cross-linking by mass spectrometry. Also, we did not attempt XL-MS with an ATP analog that lacks the primary amine, 6-methyl ATP, because this analog did not activate Mlh1-Pms1’s latent endonuclease activity.

We identified a large number of crosslinks involving the IDRs in Mlh1-Pms1 (76 of 102 crosslinks; Fig 2A and Appendix Table S3), indicating that the IDRs are highly flexible, solvent exposed, and are capable of interacting with nearby partners. Crosslinks were also seen between the Mlh1 and Pms1 C-terminal domains, consistent with the C-terminal domains being required for Mlh1-Pms1 interactions (Gueneau *et al*, 2013), and between the ATP binding (N-terminal) domains of Mlh1 and Pms1, consistent with structural and biochemical studies showing that these domains interact when both domains are ATP bound (Tran & Liskay, 2000; Ban *et al*, 1999). Interestingly, many of the 15 inter-subunit crosslinks map to regions of Mlh1 and Pms1 that are genetically critical for their function (see asterisks in Fig 2A, Appendix Table S3; Tran & Liskay, 2000; Plys *et al*, 2012; Argueso *et al*, 2003; Smith *et al*, 2013). Lastly, crosslinks were identified between the Mlh1 IDR and the Pms1 C-terminal endonuclease domain, providing evidence for models in which the condensation of the N- and C-termini of MLH proteins is linked to the activation of MLH functions (Sacho *et al*, 2008; Ban *et al*, 1999; Pillon *et al*, 2010; Gueneau *et al*, 2013).

**Figure 2.**
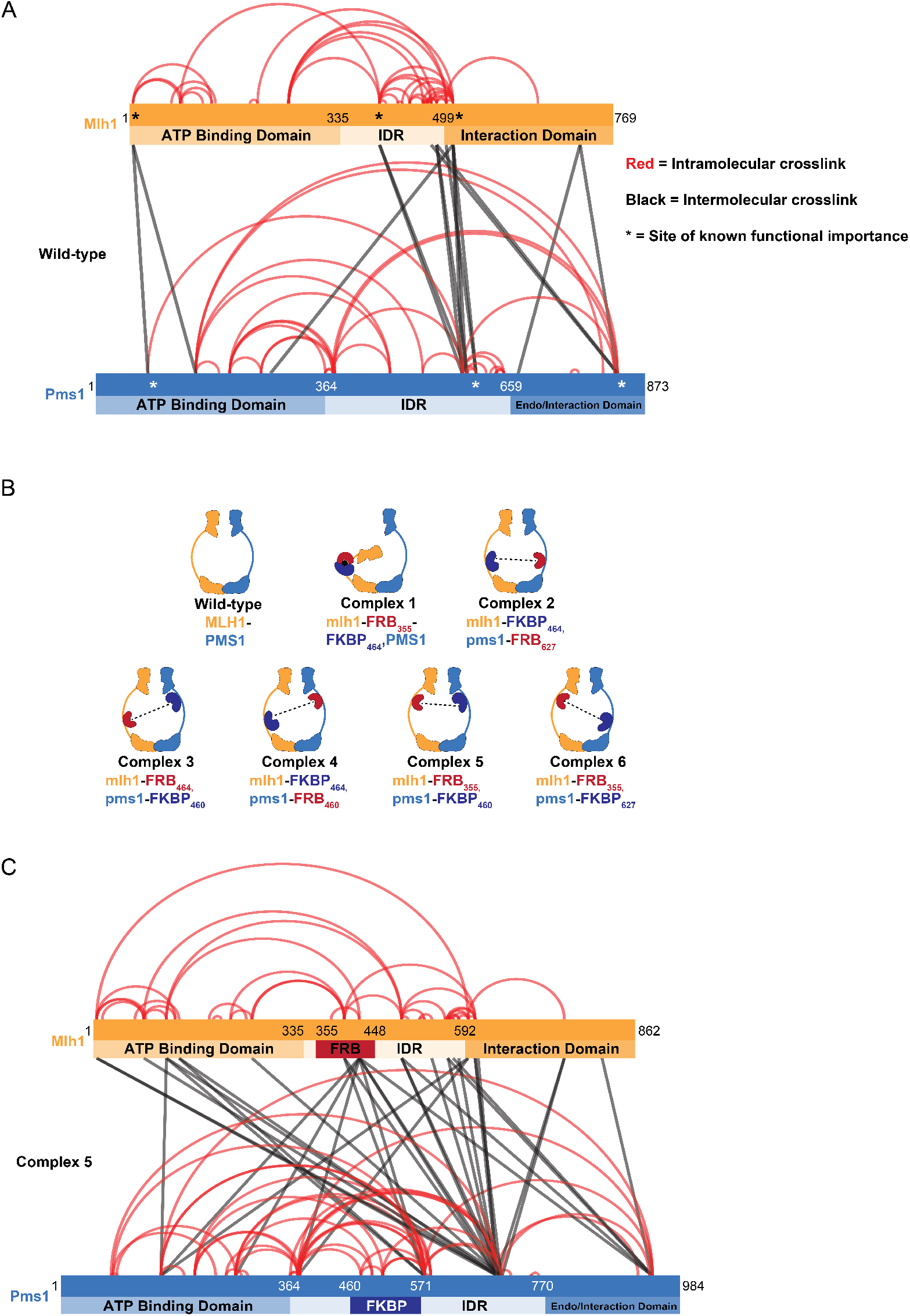
Crosslinking mass spectrometry of Mlh1-Pms1 identifies IDR regions that display limited or extensive interactions within a subunit or with the other subunits. (A) Mlh1-Pms1 was crosslinked with DSSO and then subjected to cross-linking mass spectrometry (XL-MS; Materials and Methods; Appendix Table S3). Shown are the inter- and intra-subunit lysine residue crosslinks with respect to the ATP binding, IDR, and endonuclease/interaction domains of Mlh1 and Pms1. Crosslinks involving the same position are indicated by the density of the line. * Represent locations in Mlh1 and Pms1 shown previously to disrupt MMR. (B). Complexes 1 to 6 analyzed in this study. Crosslinking analysis and previous deletion analysis of Mlh1 and Pms1 IDRs (Plys *et al*, 2012; Appendix Fig S1A) encouraged insertion of FRB and FKBP domains in the IDRs of Mlh1 and Pms1 in the indicated positions. Potential interactions with rapamycin are indicated by the dotted black lines. (C) XL-MS analysis of Complex #5 *(mlh1-FRB_355_, pms1-FKBP_460_*) with the insertions of FRB and FKBP indicated.

### Rapamycin-induced disruption of MMR *in vivo*

The identification of regions within the IDRs of Mlh1 and Pms1 that had few if any crosslinks (Fig 2A) and could be deleted without impacting MMR functions (Appendix Fig S1A; Plys *et al*, 2012) encouraged us to develop a method to restrict any potential coordinated interactions that drive MMR functions. We employed a chemically inducible system to control the dimerization of a pair of proteins with a small molecule that acts as a “dimerizer.” Dimerizers can reversibly bring regions of proteins together that normally lack interaction and have been used to study mechanistic aspects of protein localization, targeted protein degradation, transcriptional control and signal transduction (Xu *et al*, 2010; Voβ *et al*, 2015; Inobe & Nukina, 2016; Zhu *et al*, 2017; Geda *et al*, 2008). Specifically, we targeted the Mlh1 and Pms1 IDRs through the 11 kDa FKBP-rapamycin binding (FRB) domain of the mammalian target of rapamycin protein (mTOR) protein and the 12-kDa FK506 binding protein (FKBP; Fig 2A; Spencer *et al*, 1993). The FRB (93 amino acids) and FKBP (107 amino acids) polypeptides are relatively small and each bind to the small molecule macrolide rapamycin. Rapamycin binding to each domain facilitates a stable and tight FRB-FKBP heterodimer (Kd in the sub nM range; Banaszynski *et al*, 2005). We first tested if we could reversibly inhibit MMR by restricting the motion of the linker arm of Mlh1 in the presence of rapamycin (Complex 1, Fig 2B). We chose Mlh1 as our initial target because Mlh1 has been proposed to initiate Mlh1-Pms1 IDR condensation (Fig 1B; Kunkel & Erie, 2015; Sacho *et al*, 2008; Liu *et al*, 2016). We then tested the effects of disrupting the Mlh1 IDR movement genetically, using a reversion assay with a wide range of sensitivity, and biochemically, by measuring Mlh1-Pms1 DNA binding, ATPase, and endonuclease activities.

In the first experiment, FRB and FKBP were introduced in tandem into the Mlh1 IDR (Fig 2B, Appendix Fig S1A). The effects of these insertions were assessed using a highly sensitive reversion assay in which an insertion of 14 adenosine residues disrupts the *LYS2* open reading frame *(lys2-A_14_).* Frameshift mutations (primarily −1) which restore the open reading frame are detected as Lys^+^ colonies. An *mlh1Δ* strain shows a nearly four-orders of magnitude (5,000 to 6,000-fold) higher rate of reversion to Lys^+^ compared to *wild-type* (Kim *et al*, 2019; Tran *et al*, 1997). As shown in Table 1, single insertions of FRB or FKBP in Mlh1 conferred reversion rates that were only marginally higher than *wild-type* (1.8 to 4.9-fold). As shown in Fig 3A and Table 1, the double insertion allele, *mlh1-FRB_355_-FKBP_464_,* (referred to as Complex #1) conferred a very weak reversion phenotype (59-fold). The single domain insertion *mlh1* alleles conferred higher reversion rates when rapamycin was added (68 to 114-fold), but to a much lower level than seen for *mlh1Δ* (5,400-fold). One reason for the modest increase in reversion rate for the single insertion strain grown with rapamycin is that both FRB and FKBP bind rapamycin, and such binding could impact MMR by altering Mlh1 IDR flexibility. Alternatively, rapamycin binding proteins in the yeast cell other than Fpr1 (the yeast homolog of human FKBP, deleted in our strain background) may be able to weakly interact with FRB or FKBP, blocking Mlh1 from interacting with partners or mismatch substrates. Consistent with the latter idea is that the single domain insertion alleles were recessive (Table 2); if rapamycin impacted IDR function, one might have expected to still see a mutator phenotype when the FRB and FKBP insertion alleles were expressed in the presence of wild-type *MLH1,* though at lower levels (see below).

**Table 1.**
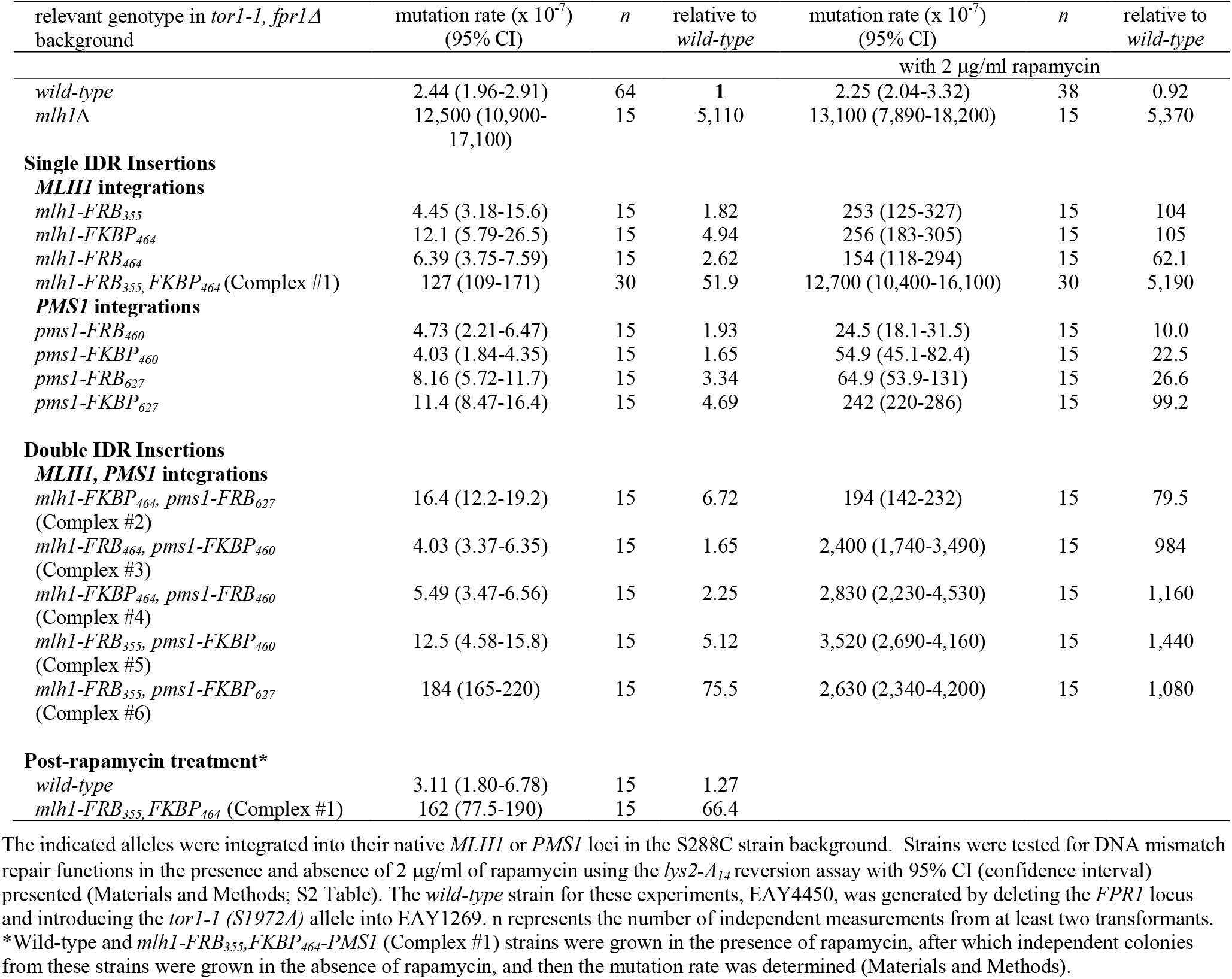
*FRB* and *FKBP* domain insertions into the *MLH1* and *PMS1* IDRs confer MMR defects in the presence of rapamycin.

**Figure 3.**
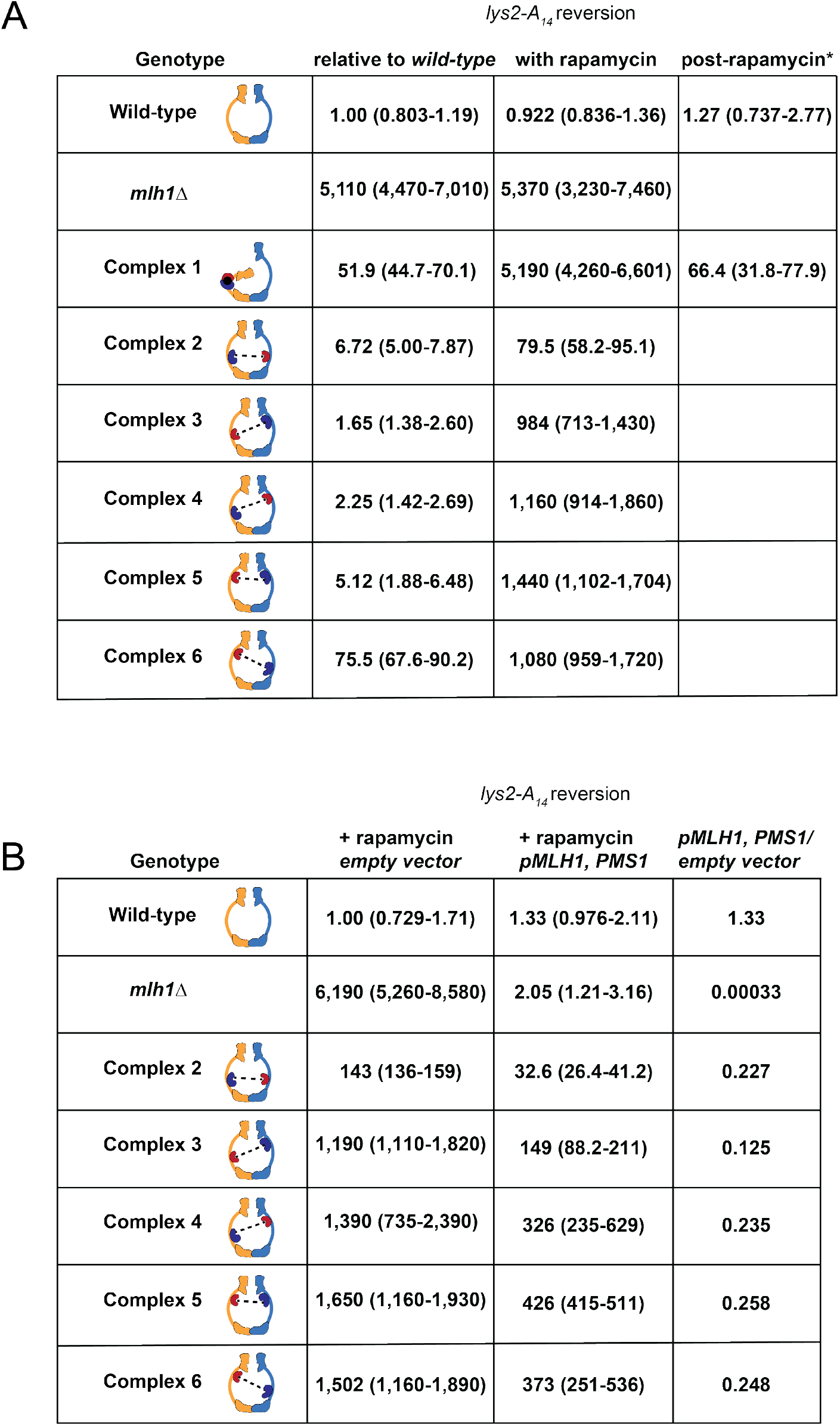
Rapamycin-induced disruption of MMR in strains expressing Mlh1-Pms1 Complexes 1-6. (A) Reversion rate analysis of strains expressing Complexes 1 to 6. Strains were grown in the absence and presence of 2 p.g/ml rapamycin and Lys^+^ reversion rates were determined using the *lys2-A_14_* assay (Materials and Methods), with a more complete data set presented in Tables 1 and 2. Data are presented relative to the rate for wild-type grown in the absence of rapamycin (2.44 × 10^-7^, Table 1) and are presented with relative 95% confidence intervals shown in parentheses. *In these experiments, independent colonies from strains grown in the presence of rapamycin were then grown in the absence of rapamycin, after which mutation rate was determined in the *lys2-A_14_* reversion assay (Materials and Methods). (B) Rapamycin induced dimerization causes a dominant MMR defect *in vivo. Wild-type* and strains containing the indicated FRB and FKBP Mlh1 and Pms1 IDR insertions, and either an empty (pRS416) or *MLH1* and *PMS1* containing *ARS-CEN* vector, were grown in the absence and presence of rapamycin (2 μg/ml) and then analyzed for reversion rate in the *lys2-A_14_* reversion assay. Detailed analysis is presented in Table 1. Data are presented relative to rate of *wild-type* containing an empty vector grown in the presence of rapamycin (2.11 × 10^-7^, Table 2). Relative 95% confidence intervals are shown in parentheses. The ratios of reversion rates for strains containing a plasmid bearing *MLH1* and *PMS1* to those containing an empty vector are shown.

**Table 2.**
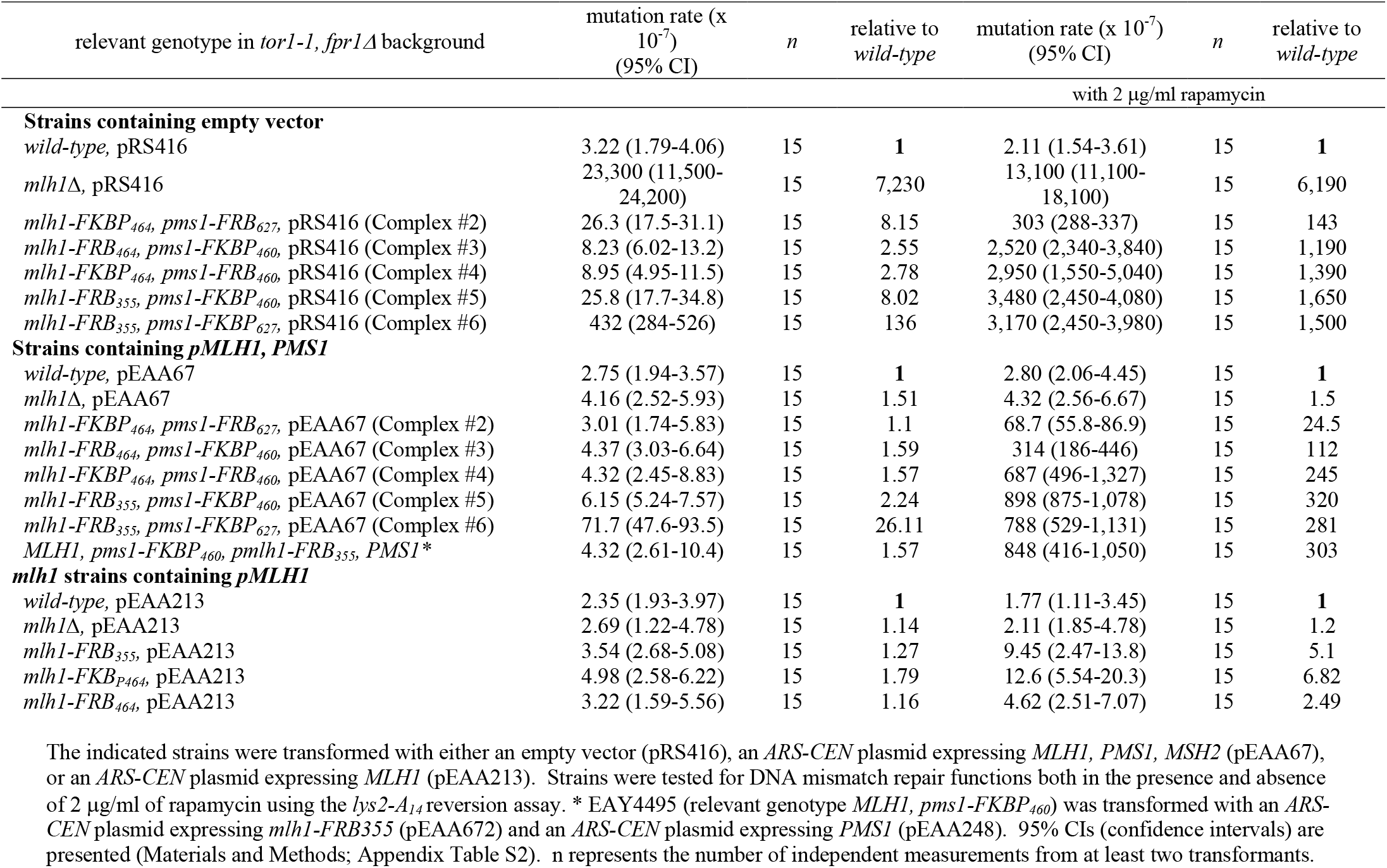
*FRB* and *FKBP* domain insertions into the IDRs of *MLH1 andPMS1* confer a dominant MMR phenotype in the presence of rapamycin.

In contrast to the single *FRB* and *FKBP mlh1* insertion strains grown in the presence of rapamycin, a complete MMR defective phenotype was observed for the *mlh1-FRB3_55_-FKBP_464_* double insertion allele (Complex #1) in the presence of rapamycin, a phenotype identical to that seen when the Mlh1 IDR was completely deleted (Plys *et al*, 2012). We then asked if MMR can be re-initiated when rapamycin is washed out of the media in *mlh1-FRB_355_-FKBP_464_* strains. With rapamycin removed, MMR activity of the *mlh1-FRB_355_-FKBP_464_* (Complex #1) strain returned to pre-rapamycin levels (Fig 3A; Table 1). This indicates that MMR functions can be reversibly and specifically modulated using a small molecule.

### Rapamycin-induced interactions between Mlh1 and Pms1 IDRs disrupt MMR

MLH clamp formation is hypothesized to create a condensed state poised for endonuclease activation upon encountering PCNA on DNA (Liu *et al*, 2016; Genschel *et al*, 2017; Kim *et al*, 2019). Based on this model we thought that restricting MLH clamp formation through FRB-FKBP interactions would inhibit MMR, and that such inhibition could be detected in biochemical assays. We constructed five additional Mlh1-Pms1 FRB/FKBP insertion complexes that maintained roughly wild-type levels of MMR in the absence of rapamycin (Fig 2B). This wildtype function was also seen when one subunit contained the FRB/FKBP insertion and the other a wild-type partner, indicating that the FRB/FKBP insertions were well tolerated (Table 2).

Complexes that restrain the IDRs of Mlh1 or Pms1 near the N-terminal region displayed similar and strong MMR defects in the presence of rapamycin (Complexes 3-6; 984 to 1,440-fold higher reversion rate to Lys^+^; Fig 3A; Table 1). Complex #2, predicted to restrain the IDRs of Mlh1 and Pms1 near their C-terminal regions, conferred weak MMR defects (79.5-fold). Interestingly, the small increase in reversion rate for Complex #2 seen in the presence of rapamycin was very similar to the rate seen when the FRB and FKBP insertion alleles in Complex #2 were expressed in the presence of a wild-type partner, indicating that the effect of rapamycin on Complex #2 function was minimal (Table 1). One explanation for these phenotypes is that interactions predicted to restrain the coordinated movements of the N-terminal ATP binding domains (Complexes 3-6) disrupt MMR whereas those predicted to maintain free movement of the N-terminal ATP binding domains (Complex #2) do not. However, we cannot exclude the possibility that the efficiency of FRB-FKBP dimerization is compromised for Complex #2.

We asked if Complexes 2-6 expressed in the presence of similar levels of wild-type complex would display evidence of subunit mixing. If this was the case, then one quarter of the complexes in a cell would be expected to contain FRB and FKBP insertions in the same heterodimer (evidence for intra-complex interactions), and the reversion rate for cells expressing both the complex and wild-type Mlh1-Pms1 in the presence of rapamycin would be expected to be one quarter of the rate seen when only the complex was expressed. If inter-complex interactions were prevalent then we would have expected to see at most a two-fold reduction in reversion rate. As shown in Fig 3B and Table 2, the reversion rates for the five complexes expressed in the presence of wild-type Mlh1-Pms1 and rapamycin were on the whole about one quarter of the rates seen when only the complexes containing FRB and FKBP domains were expressed, consistent with rapamycin inducing interactions between FRB and FKBP insertions within single heterodimers. These observations are also consistent with the determination that the local concentrations of the FRB and FKBP domains within a single heterodimer are expected to be much higher (estimated to be as high as three to four orders of magnitude) than between separate Mlh1-Pms1 molecules, which are present at ~600 copies in a single yeast nucleus (Karim *et al*, 2013). This calculation is based on a haploid yeast nucleus volume of 2.9 +/− 0.9 μm^3^ (Jorgensen *et al*, 2007) and Mlh1-Pms1 molecules in a haploid yeast nucleus diffusing randomly throughout the nuclear volume, with the motion of FRB and FKBP within the IDRs of the Mlh1-Pms1 heterodimer restricted to a shell of an approximately 10 nm radius (Gorman *et al*, 2010).

The experiments presented in Fig 3B and our calculation of Mlh1-Pms1 concentrations *in vivo* encouraged us to purify Complexes 2-5. These complexes formed stable heterodimers, remained soluble when rapamycin was added in solution, and for those tested, retained ATPase and endonuclease activities (Appendix Fig S1B and below). Because Mlh1-Pms1 forms oligomers on DNA (Hall *et al*, 2001), we were concerned that the addition of rapamycin could cause neighbouring Mlh1-Pms1 complexes to aggregate and become insoluble. However, these proteins retained solubility and activity in the presence of rapamycin, providing further evidence for rapamycin promoting intra-complex interactions.

### Rapamycin induced N-terminal dimerization prevents Complex #5 from stably binding to DNA

MLH proteins display DNA binding activities that are required for their MMR functions (Plys *et al*, 2012; Hall *et al*, 2003). To test if restriction of the IDRs of Mlh1-Pms1 affected DNA binding, we assayed binding of wild-type Mlh1-Pms1, Complex #5, and Complex #2 (Fig 4A, and Appendix Fig S2A) to a 49 bp homoduplex oligonucleotide using Microscale Thermophoresis (MST). Complexes #2 and #5 were chosen because they showed the strongest and weakest defects in MMR in cells grown in the presence of rapamycin.

**Figure 4.**
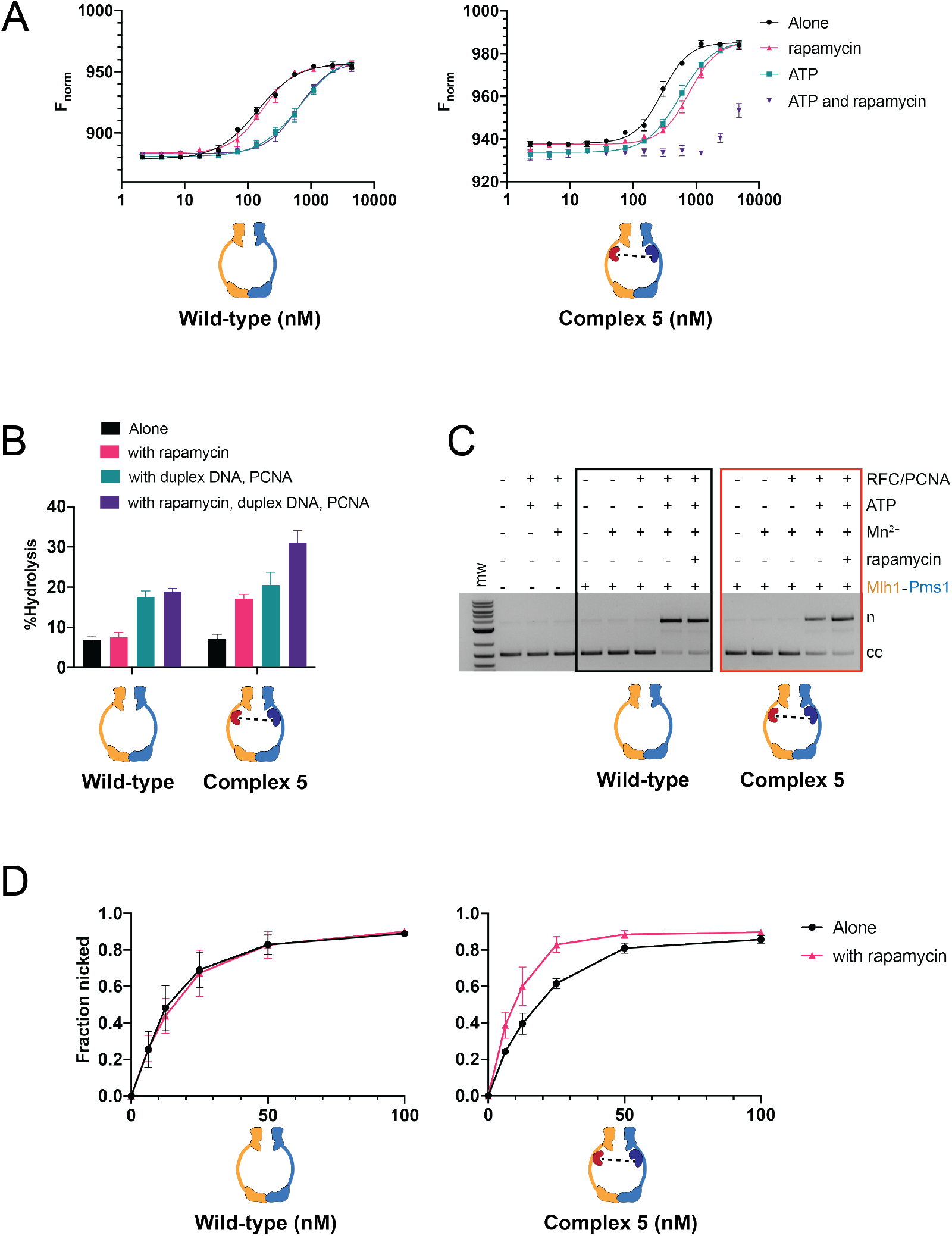
Complex #5 (mlh1-FRB_355_, pms1-FKBP_460_) displays defective DNA binding but enhanced ATP hydrolysis and endonuclease activity in the presence of rapamycin. (A) MST analysis of Mlh1-Pms1 and Complex #5 in the presence and absence of 49 bp homoduplex DNA (20 nM), ATP (1 mM), and rapamycin (1 μM). Three independent experiments (error bars indicate the mean + standard deviation) and were performed using at least two independently purified batches of each protein. F_norm_ was calculated by dividing F_hot_ (average fluorescence value in the heated state) by F_cold_ (average fluorescence value measured in the cold state before the infrared laser is turned on) and plotted as parts per thousand (%). See the Materials and Methods for details. (B) ATP hydrolysis activities of Mlh1-Pms1 and Complex #5 (0.40 μM each) were determined in the presence and absence of PCNA (0.5 μM), 49-bp homoduplex DNA (0.75 μM), PCNA (0.250 μM), and rapamycin (1 μM). Error bars indicate ± one standard deviation for three replicates. * denotes statistical significance in a student’s T-test. (C) Endonuclease activities of Mlh1-Pms1 and Complex #5 (50 nM each) determined on a closed circular DNA substrate (cc) in the presence (+) or absence (-) of MnSO_4_, ATP, rapamycin, and yeast PCNA/RFC (Materials and Methods). MnSO_4_, ATP, rapamycin, RFC and PCNA were included at 5 mM, 0.5 mM, 1 μM, 125 nM and 250 nM, respectively. n= nicked product. (D) Endonuclease activities were determined at the indicated concentrations of wild-type and Complex #5. Assays were performed in the presence of MnSO_4_, ATP, RFC, PCNA. Rapamycin was included as indicated. Error-bars indicate the standard deviation of three replicates (Appendix Fig S3).

As shown in Fig 4A and Table 3, wild-type Mlh1-Pms1 affinity for DNA was reduced in the presence of ATP (141 nM in its absence to 675 nM in its presence). This binding was unaffected by the presence of rapamycin. The reduced binding of Mlh1-Pms1 in the presence of ATP is consistent with previous studies indicating that Mlh1-Pms1 ring opening and closing is linked to interactions with DNA (Kim *et al*, 2019; Hall *et al*, 2003; Gorman *et al*, 2010). In the absence of rapamycin, Complex #5 DNA binding affinities were similar to Mlh1-Pms1 in both the presence and absence of ATP. The addition of rapamycin significantly reduced the DNA binding affinity for Complex #5 in the absence of ATP (775 nM for Complex #5 vs. 173 nM for Mlh1-Pms1). The Kd for binding of Complex #5 in the presence of ATP and rapamycin was too weak to be determined due to the aggregation of Mlh1-Pms1 at high concentrations; however, the lower limit for the Kd, based on ~50% binding, was estimated to be 5 μM compared to 628 nM for Mlh1-Pms1. In contrast, Complex #2, which showed a mild mutator phenotype in the presence of rapamycin, displayed a DNA binding activity similar to wild-type both in the presence and absence of rapamycin (Appendix Fig S2A; Table 3). Together these experiments argue that the restriction of the IDRs in Complex #5 blocks the ability of Mlh1-Pms1 to form a stable complex on DNA, thus accounting for the MMR defect seen *in vivo.* It also matches with our previous observation that a deletion of the Pms1 IDR both disrupted DNA binding and prevented ternary complex formation of MSH and MLH proteins on a mismatched substrate (Plys *et al*, 2012).

**Table 3.** K_d_ measurements (nM) for MST DNA binding analysis

|  | alone | with ATP | with rapamycin | with rapamycin and ATP |
| --- | --- | --- | --- | --- |
| Wild-type | 141 (128-157) | 675 (584-803) | 173 (157-192) | 628 (563-709) |
| Complex #2 | 207 (182-235) | 431 (380-494) | 211 (186-238) | 391 (347-444) |
| Complex #5 | 287 (264-312) | 556 (510-610) | 775 (709-855) | ND |

### N-terminal dimerization pre-activates Mlh1-Pms1 *in vitro*

To further understand why Complexes 2-6 displayed significantly increased mutation rates in the presence of rapamycin, we assayed the activity of Complex #2 and #5 in ATPase and endonuclease assays (Fig 4B; Pluciennik *et al*, 2010; Kim *et al*, 2019; Kadyrov *et al*, 2006). Previous studies showed that Mlh1-Pms1 ATPase activity is stimulated in the presence of DNA duplex oligonucleotide and PCNA (Kim *et al*, 2019; Tran & Liskay, 2000; Räschle *et al*, 2002; Genschel *et al*, 2017; Kadyrov *et al*, 2007). As shown in Appendix Fig S2B, we found that Mlh1-Pms1, Complex #2, and Complex #5 displayed, in the absence of rapamycin, similar ATPase activities. These activities were similar in both the presence or absence of DNA and PCNA. For Complex #2, rapamycin did not affect ATPase activity in the presence or absence of DNA and PCNA. However, for Complex #5 the addition of rapamycin resulted in significant increases in ATPase activities (~two-fold) both in the absence and presence of DNA and PCNA (Fig 4B).

We next measured the endonuclease activity of Complex #2 and #5 (Fig 4C, D; Appendix Fig S2C, D). In the absence of rapamycin both complexes displayed comparable levels of PCNA stimulation of endonuclease activity compared to Mlh1-Pms1. We then performed these experiments in the presence of rapamycin and found that rapamycin did not impact Mlh1-Pms1 or Complex #2 endonuclease activity; however, Complex #5 displayed an endonuclease activity that was higher (~33% elevated) than seen for Mlh1-Pms1 (Fig 4D, S2D). Taken together, these observations indicate that Complex #5 is inappropriately activated in a manner that disrupts MMR.

Finally, we analyzed Complex #5 by XL-MS in the absence of rapamycin (Fig 2C, Appendix Table S3). A similar number of crosslinks were observed for Complex #5 (132, of which 94 involved IDRs) and Mlh1-Pms1 (102 crosslinks). 34 inter-subunit crosslinks were identified for Complex #5, of which 25 were between Mlh1 and Pms1 residues. The remaining 9 were between Pms1 and FRB residues. Of the 25 inter-subunit crosslinks involving Mlh1 and Pms1 in Complex #5, 23 contained at least one residue that was involved in an inter-subunit crosslink in Mlh1-Pms1. Lastly, 6 of the 15 inter-subunit crosslinks seen in Mlh1-Pms1 were also seen in Complex #5. These observations provided evidence for conservation of inter-molecular interactions between Mlh1-Pms1 and Complex #5. Our analysis of Mlh1-Pms1 (Fig 2A) indicated that the insertions of FRB and FKBP in Complex #5 were in locations of the two proteins that were devoid of crosslinks, supporting the deletion analysis that these locations appear to not be critical for Mlh1-Pms1 function. We did not see any crosslinks in Complex #5 between FRB and FKBP, consistent with genetic data indicating that these insertions did not interact unless rapamycin was present. We were unable to perform XL-MS in the presence of rapamycin because its addition interfered with sample preparation.

Seven novel cross-links, with some identified multiple times, were observed for Complex #5 between the Mlh1 N-terminal domain and the Pms1 IDR, reminiscent of the formation of the single-arm condensed complex involving the N-terminal domain of Mlh1 observed by Sacho *et al* (2008). We also saw a shift to form crosslinks between the N-terminal and IDR domains of Mlh1 and the IDR domain of Pms1. One explanation for these changes is that the addition of new FRB/FKBP domains into the IDRs is responsible for the novel crosslinks. Alternatively, the insertions shift Complex #5 into a position where it is poised to perform the condensation steps that are normally seen in the presence of ATP. Restricting such steps, as was done in the presence of rapamycin, could thus provide an explanation for the reduced MMR functions seen for strains containing Complex #5 grown in the presence of rapamycin.

## Discussion

In this study we examined roles for the MLH IDRs in coordinating the recruitment of MLH proteins during MMR. We found that handcuffing the IDRs of Mlh1 and Pms1 through the use of rapamycin-induced FRB-FKBP dimerization disrupted MMR. This phenotype and biochemical analysis of a defective complex provided evidence that the MLH IDRs undergo coordinated intra and inter-subunit interactions (Fig 5; Kim *et al*, 2019; Hall *et al*, 2003; Gorman *et al*, 2010).

**Figure 5.**
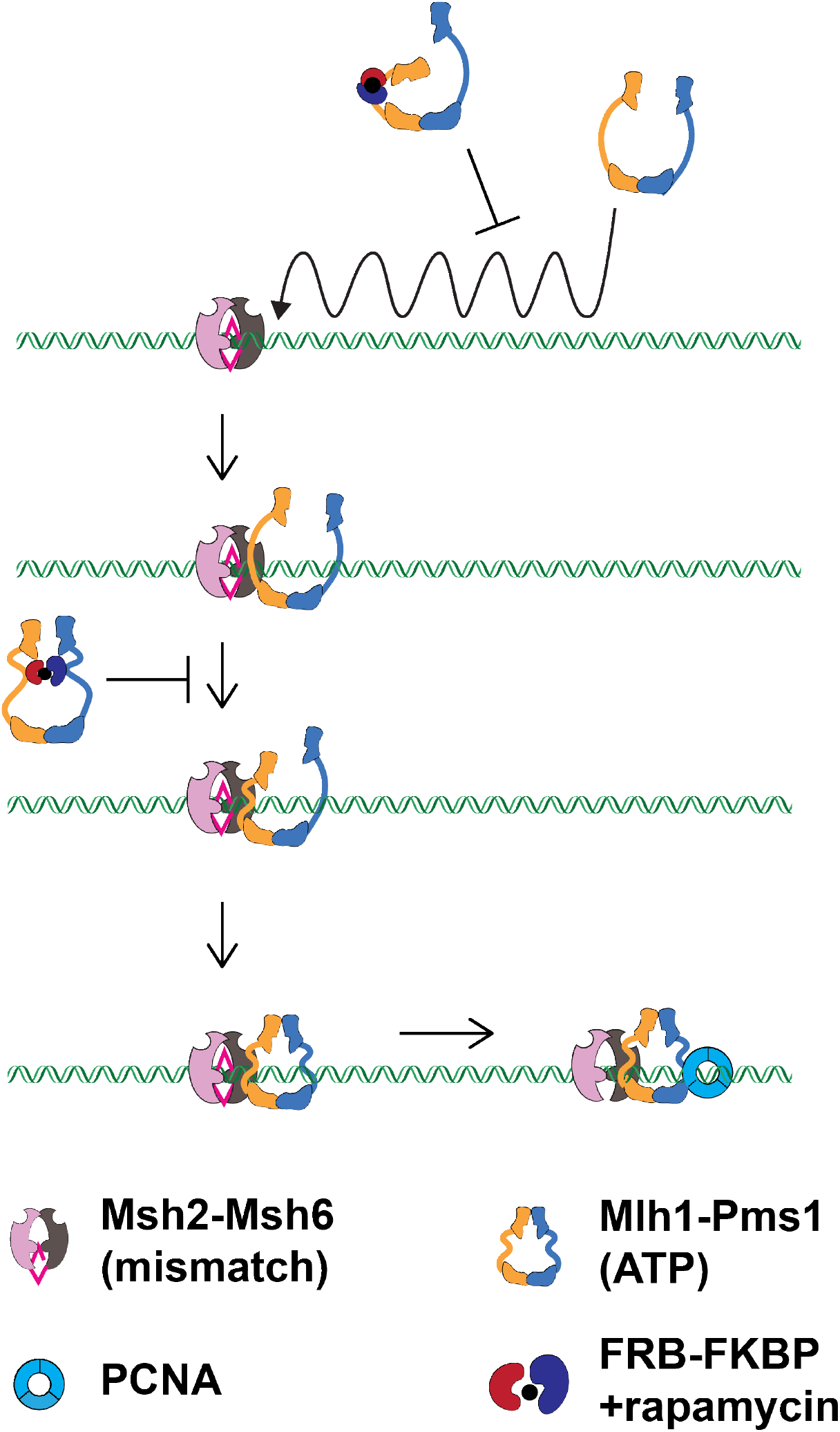
A model explaining how rapamycin induced dimerization of FRB-FKBP IDR insertions disrupts MMR. Summary of results indicating that handcuffing of the Mlh1 and Pms1 IDRs disrupts MMR by preventing the formation of complexes capable of forming a stable clamp on DNA.

Our work provides molecular evidence for Mlh1 acting as a “trigger” to initiate proteinprotein and enzymatic functions in MMR (Fig 1A). Disrupting the Mlh1 IDR via intra-subunit FRB-FKBP interactions caused a complete defect in MMR. We also saw novel cross-links for Complex #5 between the Mlh1 N-terminal domain and the Pms1 IDR, suggesting the presence of a single-arm condensed complex involving the N-terminal domain of Mlh1 (Sacho *et al*, 2008). Consistent with this observation is the finding that Complex #5 displayed an increased ATPase and nicking activity in the presence of rapamycin, suggesting that it had been activated in the absence of forming a stable complex on DNA. The idea of Mlh1 serving as a trigger is intriguing because Mlh1 serves as a common subunit for three MutL complexes in baker’s yeast, Mlh1-Pms1, Mlh1-Mlh3 and Mlh1-Mlh2, where Mlh1-Mlh3 plays a minor role in MMR and a major role in the resolution of double-Holliday junctions in meiosis to form crossovers, and Mlh1-Mlh2 regulates gene conversion tract lengths in meiosis (reviewed in Furman *et al*, 2021).

In such a model, recruitment of various MLHs is accomplished through the initial interactions of Mlh1 with the particular repair pathway, followed by clamp formation by the specificity subunit (Mlh2, Mlh3, or Pms1) that activates the complex for its specific role. Thus, our studies provide mechanistic support for the idea that IDRs license Mlh1-Pms1 to interact stably interact with DNA (Fig 5) during MMR. It complements previous studies showing that IDRs regulate how a DNA repair enzyme scans chromatin for DNA lesions and how repair functions (for example, nicking of the newly replicated strand during MMR) are activated (Gorman *et al*, 2012; Kim *et al*, 2019; Gorman *et al*, 2010; Liu *et al*, 2016; Mardenborough *et al*, 2019).

### Modulation of MMR using a small molecule

We used an inducible dimerization approach to modulate MMR functions through the addition and removal of the small molecule rapamycin (Spencer *et al*, 1993). The reversibility of this method can provide a valuable means to understand how an increased reversion rate can provide beneficial mutations to an organism adapting to changing environments (reviewed in Raghavan *et al*, 2018; 2019). Elevated mutation rates, while providing a source of beneficial mutations for adaptation, are associated with long-term fitness costs due to the accumulation of deleterious mutations (Lynch *et al*, 2016; Bui *et al*, 2015, 2017). In the *lys2-A_14_* reversion assay where the difference between *wild-type* and MMR defective is roughly 5,000-fold, we identified complexes that showed *wild-type* or near *wild-type* phenotypes in the absence of rapamycin, but a wide-range of elevated reversion rates in its presence (Fig 3A). For example, in the presence of rapamycin, Complex #2 conferred a 79.5-fold elevated rate compared to *wild-type,* Complex #3, a 984-fold higher rate, Complex #5, a 1,440-fold higher rate, and Complex #1, a rate indistinguishable from *mlh1* null (5,190-fold). Most informative was that the null phenotype in Complex #1 could be reversed to the level observed in the absence of rapamycin. This resource can thus allow one to grow cells in stressed environments (such as high salt) and then pulse them with rapamycin for different time intervals under a variety of mutation supply conditions. Such experiments would allow one to determine the impact of cellular fitness as a function of mutational load and would be relevant to understand the progression of disease states accelerated by increased mutation rate.

Mlh1-Mlh3 has been shown in meiosis in both lower and higher eukaryotes to facilitate the resolution of double-Holliday junction intermediates into crossovers. These crossovers facilitate the segregation of homologs in the first meiotic division (reviewed in Manhart & Alani, 2016). The use of a small molecule to potentially modulate the MMR and crossover resolution functions of Mlh1-Mlh3 at different stages in meiosis could thus be valuable in teasing apart proposed separable functions for MLH proteins in meiosis (Claeys Bouuaert & Keeney, 2017).

### An application of the FRB-FKBP dimerization system

IDRs are characterized as being highly dynamic and without well-defined structures. However, many IDRs have been shown to undergo disorder to order transitions, and in some cases form different structures when complexed with different partners (reviewed in Van Der Lee *et al*, 2014; Uversky, 2019). Interestingly a length distribution analysis of human proteins obtained from the SwissProt data base indicated that very long IDRs (>500 amino acids) were found in higher numbers than predicted (Tompa & Kalmar, 2010). Furthermore, gene ontology analysis indicated that IDRs greater than 500 amino acids in length are overrepresented in transcription functions and those between 300 and 500 amino acids in length are enriched in kinase and phosphatase functions (Lobley *et al*, 2007). Thus, our strategy to introduce FRB and FKBP domains into the IDRs of the MLH proteins could be applied to reversibly block or lock in specific conformational states of IDRs that regulate a variety of cellular processes.

## Materials and Methods

### Media

*S. cerevisiae* strains were grown at 30°C in either yeast extract-peptone (YPD) media or minimal selective media (SC; Rose *et al*, 1990). When required, geneticin (Invitrogen, San Diego) was added at 200 μg/ml (Goldstein & McCusker, 1999).

### Strains

S288c background derived yeast strains are listed in Appendix Table S1, with details regarding their construction available upon request. Briefly, the rapamycin resistant strain EAY4450, derived from EAY1269, was constructed as described by Zhu *et al* (2017) for the yeast strain JJY70. Plasmids bearing *MLH1* and *PMS1* derivatives (Appendix Table S2) were digested with the appropriate restriction enzymes and introduced into EAY4450 using methods described in Rose *et al* (1990) and Gietz *et al* (1995). *mlh1Δ* derivatives of EAY4450, EAY4488-4490, were constructed by digesting pEAI160 *(mlh1Δ::KanMX)* with the *SphI* and *KpnI* prior to transformation. All integrants were genotyped by PCR using primers that map outside of the restriction sites used for integration, and the presence of specific alleles was confirmed by DNA sequencing.

### Plasmids

Plasmids used in this study are listed in Appendix Table S2. Full details of plasmid constructions and maps are available upon request. Briefly, integration vectors (pEAA672, 674, 675, 713 derived from pEAA213-*MLH1::KanMX;* pEAI453, 454, 455, 468, derived from pEAA238-*PMS1*) containing *FRB* or *FKBP* insertions were constructed using HiFi Gibson cloning (New England Biolabs, Ipswich, MA), with PCR fragments generated from pEAA213 or pEAA238, and gBlocks (Integrated DNA Technologies, Coralville, IA) encoding FRB and FKBP protein domains (Plys *et al*, 2012; Xu *et al*, 2010; Geda *et al*, 2008). The FRB and FKBP domains were inserted immediately after the indicated amino acid position of Mlh1 and Pms1 (Appendix Fig S1A). *PMS1* integration vectors were constructed through HiFi Gibson cloning by inserting the *LEU2* gene from pRS415 (Christianson *et al*, 1992) downstream of *PMS1* in the *ARS-CEN* vectors pEAA671, 676, 677, and 678. *MLH1* (pEAE 269, 446, 447, 448) and *PMS1* (pEAE431, 433, 435) protein expression vectors were derived from pMH1 *(GAL1-MLH1-VMA-CBD,2μ, TRP1)* and pMH8 *(GAL10-PMS1,2μ, LEU2),* respectively (Hall & Kunkel, 2001). The DNA sequence of the open reading frames (including 300 bp upstream and 150 bp downstream) of constructs was confirmed by Sanger DNA sequencing (Cornell BioResource Center). The amino acid sequence of the FRB insertions, with glycine (G) and serine (S) linkers shown in bold is: **GS**ILWHEMWHEGLEEASRLYFGERNVKGMFEVLEPLHAMMERGPQTLKETSFNQAYGRDLM EAQEWCRKYMKSGNVKDLLQAWDLYYHVFRRISK**GS**. The amino acid sequence of the FKBP insertions, with glycine and serine linkers shown in bold is: **GS**GVQVETISPGDGRTFPKRGQTCVVHYTGMLEDGKKFDSSRDRNKPFKFMLGKQEVIRGWE EGVAQMSVGQRAKLTISPDYAYGATGHPGIIPPHATLVFDVELLKLE**GS** (Spencer *et al*, 1993; Geda *et al*, 2008).

### *lys2::insE-A_14_* reversion assay (Tables 1, 2)

Assays were performed as described previously (Kim *et al*, 2019; Kim *et al*, 2019). Briefly, strains listed in Appendix Table S1 were freshly struck from frozen stocks and grown in synthetic complete media in the presence or absence of rapamycin (2 μg/ml) and then inoculated in liquid complete media maintaining either the presence or absence of rapamycin prior to plating onto complete and lysine dropout plates. Strains containing plasmids were grown in minimal selective leucine dropout, to maintain pEAA213, or uracil dropout, to maintain pRS416 and pEAA67. Rapamycin was included in growth media until just before cell cultures were plated onto complete and lysine dropout plates to measure *lys2::insE-A_14_* reversion. Rates of *lys2::insE-A_14_* reversion were calculated as μ=f/ln(N·μ), where f is reversion frequency and N is the total number of revertants in the culture (Tran *et al*, 1997). For each strain, 15-64 independent cultures, obtained from two to three independent transformants bearing a unique allele, were assayed on at least two different days to prevent batch effects, and 95% confidence intervals and all computer-aided rate calculations were performed as described previously (Kim *et al*, 2019; Dixon & Massey, 1969). For the post-rapamycin treatment described in Table 1, the *wild-type* and Complex #1 strains were grown both in the presence and absence of rapamycin (2 μg/ml) and then inoculated in liquid complete media maintaining either the presence or absence of rapamycin prior to plating onto complete and lysine dropout plates. Upon colony counting after 3 days of incubation, a single colony from the complete plate was inoculated in liquid media lacking rapamycin and then assayed again for mutator phenotype.

### Cross-linking Mass Spectrometry (XL-MS)

Mlh1-Pms1 and Complex #5 were crosslinked with disuccinimidyl sulfoxide (DSSO; Thermo Fisher Scientific). A 50 mM stock solution of DSSO was freshly prepared by dissolving DSSO into anhydrous dimethyl sulfoxide (DMSO). 5 μg of Mlh1-Pms1 and Complex #2 and #5 derivatives were incubated in 50 μl of buffer containing 25 mM HEPES pH 8.0, 180 mM NaCl, 10% glycerol). DSSO was included at a final concentration of 1.25 mM and the reaction was then incubated for 30 min at room temperature. Reactions were quenched by the addition of Tris-HCl pH 8.0 to a final concentration of 10 mM. Samples were digested and processed for MS as described in Yugandhar *et al* (2020). In brief, the cross-linked samples were denatured with 1% sodium dodecyl sulfate (SDS) at 65 °C for 15 min, reduced by 1 mM dithiothreitol (DTT) at room temperature for 15 min, and then alkylated with 25 mM iodoacetamide at room temperature for 15 min. Proteins were precipitated by adding 3X volumes of cold acetone/ethanol/acetic acid solution (50:49.9:0.1, v/v/v). The precipitates were resuspended in 8 M Urea, 50 mM Tris-Cl and 150 mM NaCl, pH 8.0. After dilution to 2 M Urea, trypsin (Trypsin Gold, Promega, Madison, WI) digestion was performed at 37 °C overnight. Trifluoroacetic acid-formic acid (TFA-FA) solution was applied to terminate digestion. The digested peptides were desalted using Sep-Pak C18 cartridge (Waters, Dublin, Ireland), dried using SpeedVac™ Concentrator (Thermo Fisher Scientific, Pittsburgh, PA) and stored in −80 °C for further analysis.

Peptides were resuspended in 0.1% TFA and then analyzed using an Orbitrap Fusion Lumos Tribrid mass spectrometer (Thermo Fisher Scientific) coupled online to an EASY-nLC 1200 system (Thermo Fisher Scientific) equipped with a 75-μm × 25-cm capillary column (inhouse packed with 3-μm C18 resin; Michrom BioResources). For each analysis, peptides were eluted using a 45 min liquid chromatography gradient (5% B-40% B; mobile phase A composed of 0.1% formic acid (FA), and mobile phase B composed of 0.1% FA and 80% acetonitrile (ACN)) at a flow rate of 200 nl/min.

The CID-MS2-HCD-MS3 acquisition method was used for DSSO-crosslinking identification. The MS1 precursors were detected in the Orbitrap mass analyzer at resolution of 60,000 with a scan range from 375 m/z to 1500 m/z. The precursor ions with the charge of +4 to +8 were selected for MS2 analysis at a resolution of 30,000 (AGC target = 1 × 10^5^, precursor isolation width = 1.6 m/z, and maximum injection time = 10^5^ ms), with the normalized collision energy of CID at 25%. The two most abundant reporter ions with a mass difference of 31.9721 Da in CID-MS2 spectra were selected for further MS3 analysis. The selected ions were fragmented in an Ion Trap using HCD with the normalized collision energy at 35% and AGC target of 2 × 10^4^. All spectra were recorded by Xcalibur 4.1 software and Orbitrap Fusion Lumos Tune Application v. 3.0 (Thermo Fisher Scientific).

The cross-link search was performed using MaXLinker software (Yugandhar *et al*, 2020). The wildtype and engineered Mlh1-Pms1 cross-linked samples were searched against database with corresponding target and randomized sequences, along with randomized sequences of *E. coli* proteome (5268 sequences) for efficient FDR estimation. Crosslink maps for Mlh1-Pms1 are a composite of results from two independent crosslink trials.

### Protein purification

Mlh1-Pms1 and Complex #2 to #5 variants (Appendix Fig S1B) were purified as described from galactose-induced cultures of BJ2168 *(MATa, ura3-52, leu2-3,112, trp1-289, prb1-1122, prc1-407, pep4-3)* bearing pEAE expression vectors (Appendix Tables S1, S2; Plys *et al*, 2012; Hall & Kunkel, 2001). The *MLH1* pEAE672, 674, 675 and 713 expression constructs contain a FLAG tag at position 499 (with respect to the wild-type sequence). RFC and PCNA were expressed and purified from *E. coli* (Cooper *et al*, 1999; Finkelstein *et al*, 2003).

### DNA substrates for biochemical assays

Closed circular pUC18 (2.7 kb, Thermo Fisher Scientific, Waltham, MA) was used as the DNA substrate in the endonuclease assays presented in Fig 4, Appendix Fig S2. A 49-mer homoduplex DNA substrate was included in the ATPase experiments presented in Fig 4B and S2B. This substrate was made by annealing AO3142 (5’GGGTCAACGTGGGCAAAGATGTCCTAGCAAGTCAGAATTCGGTAGCGTG) and AO3144 (5’CACGCTACCGAATTCTGACTTGCTAGGACATCTTTGCCCACGTTGACCC). AO3142 and AO3144 were added at an equal molar ratio in buffer containing 10 mM Tris-HCl, pH 7.5, 100 mM NaCl, 10 mM MgCl_2_, and 0.1 mM EDTA. These oligonucleotides were annealed through an incubation at 95 °C for 5 min, followed by cooling to 25 °C at a rate of 1 °C/min. Following annealing, excess single-stranded DNA was removed using an S300 spin column (GE Healthcare). For the microscale thermophoresis (MST) analysis presented in Fig 4A, Appendix Fig S2A, a 48-mer homoduplex DNA substrate was made by annealing AO4549 (5’-6-FAM-CTGGACGGGTTAAGACCGAACGTGGCTCCAGAAACGGGTGCAACTGGG; synthesized by Integrated DNA technologies) and AO4548 (5’CCCAGTTGCACCCGTTTCTGGAGCCACGTTCGGTCTTAACCCGTCCAG) as described above.

### Endonuclease assays

Reactions were performed in 20 μl in 1X endonuclease buffer (20 mM HEPES-KOH (pH 7.5), 20 mM KCl, 2.5 mM MnSO_4_, 0.2 mg/mL BSA, 1 % glycerol; Rogacheva *et al*, 2014). Mlh1-Pms1, Complex #2, or Complex #5 were included at final concentrations of 6.25 to 100 nM. When indicated, rapamycin was included at a final concentration of 1 μM. Rapamycin was dissolved into DMSO at a concentration of 10 mM then serially diluted in DMSO to a final concentration of 100 μM. The 100 μM solution was then diluted in 1x endonuclease reaction buffer to a concentration of 20 μM before being added to individual reaction tubes at a final concentration of 1 μM. Rapamycin was added to the reaction before the addition of the DNA substrate and the reaction was incubated at 37°C for 5 mins prior to DNA addition. Reactions (37°C, 40 min) were started following the addition of pUC18 (5.1 nM final concentration of plasmid) and stopped by the addition (final concentrations shown) of 0.1 % SDS, 14 mM EDTA, and 0.1 mg/ml Proteinase K (New England Biolabs). DNA was electrophoresed in 1.2% agarose gels in 1xTAE (Tris-acetate-EDTA) containing 0.1μg/mL^-1^ ethidium bromide, which causes covalently closed circular DNA isoforms to separate from nicked DNA product. Gels were run in 1x TAE at 100 V for 45 min. Negative control lanes were used as background and were subtracted out of reported quantifications. BioRad Image Lab Software, v5.2.1 was used to quantify gels.

#### ATPase assays

ATPase activity was determined using the Norit A absorption method described previously (Kim *et al*, 2019; Rogacheva *et al*, 2014). Briefly, 30 μl reactions contained 0.4 μM of Mlh1-Pms1, Complex #2, or Complex #5, 100 μM [γ-32P]-ATP, 20 mM Tris, pH 7.5, 2.0 mM MgCl_2_, 0.1 μM DTT, 1 mM MnSO_4_, 75 mM NaCl, 1% glycerol, 40 μg/ml BSA. Reactions were incubated for 40 min at 37 °C. When specified, DNA (49-mer homoduplex DNA substrate), PCNA, and rapamycin were included at 0.75 μM, 0.5 μM, and 1 μM, respectively. Just prior to addition, rapamycin was diluted and included in reactions as described for the endonuclease assays.

### Microscale Thermophoresis (MST) assay

MST was performed using a Monolith NT.115 instrument (NanoTemper Technologies) equipped with red and blue filters using methods described in Hosford *et al*, 2020). MLH complexes were serially diluted 16 times in two-fold steps to final concentrations of 4,400 to 2.14 nM for Mlh1-Pms1,4,000 to 1.95 nM for Complex #2, and 4,800 to 2.34 nM for Complex #5, and then mixed with a stock of 6-FAM-labeled duplex oligonucleotide (40 nM, sequence described above). Reaction buffer contained 20 mM Tris-HCl (pH 7.5), 20 mM NaCl, 0.01 mM EDTA, 2 mM MgCl_2_, 40 μg/mL^-1^ BSA, 0.1 mM DTT and 0.05% TWEEN-20. Assays with nucleotide, rapamycin, or both contained 1 mM ATP and 1 μM rapamycin. ATP and rapamycin were added to reactions containing Mlh1-Pms1 prior to DNA addition. The reactions (20 μl volumes) were then incubated for 5 min at room temperature. Following DNA addition, reactions were incubated at room temperature at 30 °C for 15 min. They were then loaded into standard capillaries (NanoTemper Technologies). Reactions were then tested with 40% excitation power, medium MST power, and measured using M.O. Control software (NanoTemper Technologies). M.O. Affinity Analysis software (version 2.3, Nanotemper Technologies) was used to analyze data and determine the normalized fluorescence (F_nom_) for each concentration. F_norm_ is calculated by dividing F_hot_ (average fluorescence value in the heated state) by F_cold_ (average fluorescence value measured in the cold state before the infrared (IR) laser is turned on) and plotted as parts per thousand (%). Three independent reactions were measured to obtain F_norm_ values, which were then averaged (mean ± standard deviation) and plotted against the respective concentration of Mlh1-Pms1 (Fig 4A). Binding constants (K_d_) were determined by nonlinear curve fitting using GraphPad Prism 9. All experiments were performed using at least two independently purified proteins.

## Acknowledgments

We are grateful to Eric Greene for suggesting the use of the FRB-FKBP dimerization to probe Mlh1-Pms1 functions, and Ilya Finkelstein, Carol Manhart, Joshua Chappie, Philip Cole, Sy Redding, Myfanwy Adams, Fred Dyda, Alison Hickman, and Jeff Jorgensen for providing their expertise, advice, and use of equipment. C.M.F. and E.A. were supported by the National Institute of General Medical Sciences of the National Institutes of Health (https://www.nih.gov/): R35GM134872, and T.-Y.W., Q.Z., K.Y., and H.Y. by R01GM124559. The content of this study is solely the responsibility of the authors and does not necessarily represent the official views of the National Institutes of Health. The funders had no role in study design, data collection and analysis, decision to publish, or preparation of the manuscript.

## Author contributions

CMF performed the experiments described in Figures 3, 4, S1, S2 and S3, and Tables 1–3. CMF performed the DSSO cross linking for the MLH complexes presented in Figure 2 and Table S3, with the mass-spectrometry analyses performed by T-YW, QZ, and KY. CMF, T-YW, QZ and EA wrote the initial draft of the manuscript and all authors participated in preparing and revising the manuscript.

## Data Availability Section

This study includes no data deposited in external repositories.

